# TmDOTP : An NMR- based Thermometer for Magic Angle Spinning NMR Experiments

**DOI:** 10.1101/566729

**Authors:** Dongyu Zhang, Boris Itin, Ann E. McDermott

## Abstract

Solid state NMR is a powerful tool to probe membrane protein structure and motions in native lipid structures. Sample heating, caused by magic angle spinning and radio frequency irradiation in solid state NMR, produces uncertainties in sample temperature and thermal broadening caused by temperature distributions, which can also lead to sample deterioration. To measure the sample temperature in real time, and to quantify thermal gradients and their dependence on radio frequency irradiation or spinning frequency, we use the chemical shift thermometer TmDOTP, a lanthanide complex. Compared to other NMR thermometers (e.g., the proton NMR signal of water), the proton spectrum of TmDOTP exhibits higher thermal sensitivity and resolution. In addition, the H_6_ proton in TmDOTP has a large chemical shift (−175 ppm at 275 K) and is well resolved from the rest of the proton spectrum. We identified two populations of TmDOTP, with differing temperatures and dependency on the radio frequency irradiation power, within proteoliposome samples. We interpret these populations as arising from the supernatant and the pellet, which is sedimented from the sample spinning. Our results indicate that TmDOTP is an excellent internal standard for monitoring temperatures of biophysically relevant samples without distorting their properties.

## Introduction

Magic angle spinning (MAS) solid-state nuclear magnetic resonance (SSNMR) is a powerful technique for studying biomolecules^1^, including: protein assemblies in near native conditions^2^1, membrane proteins^3–5^ and amyloid fibrils^6–8^. SSNMR provides rich information on protein molecular structure and motions. Restricted global molecular motions in solids allow for the retention of dipolar couples, which enables direct measurement of distances and local orientations. Many of the most exciting developments in SSNMR however, involve pulse sequences with long and strong radio frequency (RF) irradiation elements; Sample heating from magic angle spinning and RF irradiation has been cause for concern^9,10^ Elevated and uncalibrated temperatures within the sample complicates the interpretation of dynamics and other properties. Moreover, heating gradients within the MAS rotor may contribute to peak broadening.

Sample heating originates in part from friction between bearing gas and the rotor during MAS^10,11^. Heat is also generated from RF irradiation during high power decoupling due to inductive dielectric heating on conductive or dipolar samples^9,12,13^. The application of high power oscillating electric field causes free charges and permanent electric dipoles to move, generating kinetic energy, which dissipates in the surrounding sample as heat^14^. The absorption of RF energy is maximized when ωτ = 1, where ω represents the frequency of oscillating field and τ is the characteristic relaxation time of the molecule^14,15^. RF heating is of particular concern in SSNMR experiments on biological samples due to the resistive losses from the high concentration of ions in typical biological buffers and the dipolar losses from the presence of mobile permanent dipoles such as in hydrated lipids^12,15–18^. Since the heating mechanism is difficult to completely avoid for such samples, it is critical to monitor the temperature changes and heating gradient in order to control the sample temperature during SSNMR experiments.

Here, we use thulium 1,4,7,10-tetra-azacyclododecane-l,4,7,10-tetrakis (methylene phosphonate) TmDOTP^5-^ (CAS: 30859-88-8), specifically the H_6_ proton chemical shift, as an internal thermometer to measure the temperatures for biological samples during SSNMR experiments (Figure 1A). TmDOTP^5-^ is a water soluble paramagnetic complex, which is known to have strongly temperature dependent chemical shifts for ^1^H, ^13^C and ^31^P (Figure 1B). Compared with other compounds that have excellent thermal resolution in chemical shifts, such as Pb(NO_3_)_2_, KBr or Sm_2_Sn_2_O_7_^19^, TmDOTP^5-^ is convenient to measure, since temperature measurements are made *in situ*, without changing samples or probe tuning. The H_6_ proton was chosen for its moderately high temperature sensitivity and relatively narrower linewidth compared to the other five nonequivalent protons^20^. Due to its low toxicity, TmDOTP^5-^ has been applied to clinical magnetic resonance to measure the temperature of tissue cells and tumor cells during surgery^20^.

**Figure 1.**
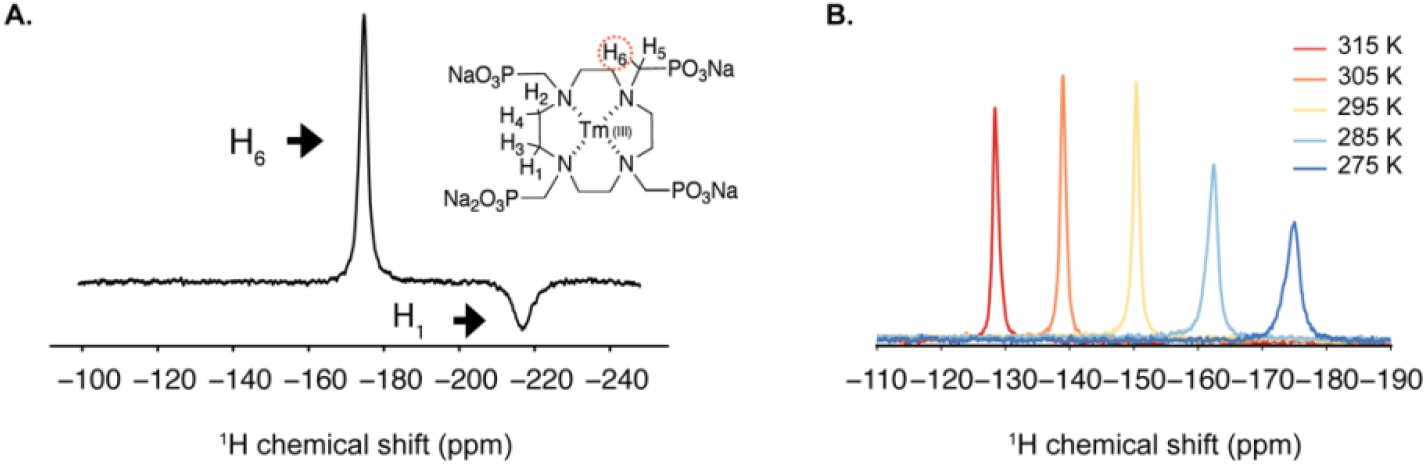
(A) Molecular structure of TmDOTP with H_6_ highlighted and portion of the ^1^H NMR spectrum of 25mM TmDOTP. The sample contains KcsA proteoliposome and 25mM TmDOTP. H_6_ is at −175 ppm while H_1_ is −217 ppm. (B) Overlay of the spectra of the H_6_ proton in TmDOTP acquired at various temperatures. All spectra were collected on 900 MHz with MAS frequency at 5 kHz. Chemical shift was referenced to the TMS at 0 ppm.

## Experiment and Method

### Sample preparation

TmDOTP (Macrocyclics, Inc.) buffer was made with 25mM TmDOTP (molecular weight: 914.2 g/mol), 20 mM MOPS and 100 mM KCl at pH 7.5 in 99.96% D_2_O (Sigma). 10mg wt-KcsA was overexpressed and reconstituted into 3:1 DOPE/DOPG (wt/wt) liposomes as described previously^21^. Then the proteolipsome sample was resuspended and incubated with TmDOTP buffer (same conditions as above) for 2 hours before packing into a regular-wall zirconia Bruker 3.2mm rotor with a silicon spacer on the top.

### NMR Spectroscopy

Experiments were carried out using a 3.2 mm standard-bore E-free probe and 1.3 mm HCN probe on a Bruker Avance II 900 MHz spectrometer. The temperature was regulated with VT gas (flow rate 1070 L/hr) and a heater in the probe. The VT control unit was calibrated using the chemical shift difference between the -CH_3_ and -OH groups of methanol^22^. The temperature of the system was allowed to equilibrate for at least 15 minutes after each temperature change. The pulse sequence used to measure heating from RF irradiation is shown in Figure 2. *τ*_1_ represents the duration of the heating pulse, which resembles high power decoupling. Unless otherwise specified, *τ*_1_ was kept at 30 ms and the recycle delay was 1 s. *τ*_2_ is the delay time to study cooling. Owing to the short T_1_ of the TmDOTP H_6_ proton (~ 800 μs), we kept *τ*_2_ at 5ms to limit heat dissipation before acquisition. A short spin echo (*τ*_3_ = 40 μs) is added before acquisition to suppress TmDOTP H1 signal, which has a larger temperature slope (ppm/K) and could interfere with H_6_ signal at high temperature. The one-dimensional MAS spectra were acquired using 8 dummy scans and 512 scans.

**Figure 2.**
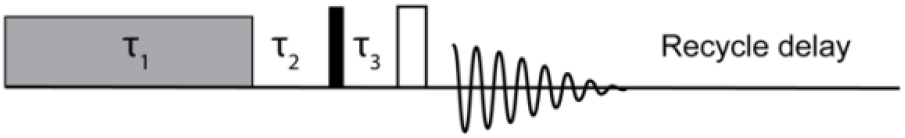
^1^H Pulse sequence used to measure the temperature increase from RF irradiation. The RF irradiation is applied for the duration of *τ*_1_ · *τ*_2_ represents the delay before proton 90 pulse. A spin echo with *τ*_3_ =20μs is applied before acquisition to dephase the signal from H_1_ in TmDOTP

The ^13^C-^13^C dipolar assisted rotational resonance (DARR^23^) experiments with 50 ms mixing time were performed on the same 900 MHz spectrometer with a MAS rate of 16.666 kHz and a set temperature of 267 K. Proton decoupling with the SPINAL64^24^ scheme at 90 kHz was applied during acquisition. The recycle delay was 2.5 s. The ^1^H and ^13^C Dual-Receiver DARR experiment was performed on 3.2 mm standard-bore E-free probe with Bruker Avance NEO spectrometer operating at 700 MHz. The MAS rate was 12.5 kHz and VT gas flow was 2000 1/h. SPINAL64 decoupling was applied at ω_1_/2π = 90 kHz on the proton channel during acquisition (15 ms) and the recycle delay was 2 s.

## Results and Discussion

### ^1^H NMR signals of H_2_O and TmDOTP as precise temperature measures

The water chemical shift is known to be sensitive to temperature and has been employed as an internal thermometer in several studies^12,18,23^. We compared the temperature dependence of the chemical shift of the H_6_ proton in TmDOTP versus the water proton in the same sample (Figure 3). The temperature dependency of the H_6_ proton in TmDOTP, 1.06 ±0.04 ppm/K, is 2 orders of magnitude larger than that of water (−1.1×10^-2^±0.1×10^-2^ ppm/K), while the full width at half maximum (FWHM), (1.5±0.6 ppm), is one order of magnitude larger than that of water (0.12±0.01 ppm). The uncertainty in the calculated temperature dependencies were dominated by the fitting error. Overall, TmDOTP allows for more accurate and precise temperature measurements than water. The homogeneous linewidth of the H_6_ proton calculated from 1/πT_2_ was about 980 ± 60 Hz at 275 K. The offset between homogeneous linewidth and the actual linewidth, 1.4 kHz, presumably is due to inhomogeneous broadening that cannot be refocused by a Hahn spin echo

**Figure 3.**
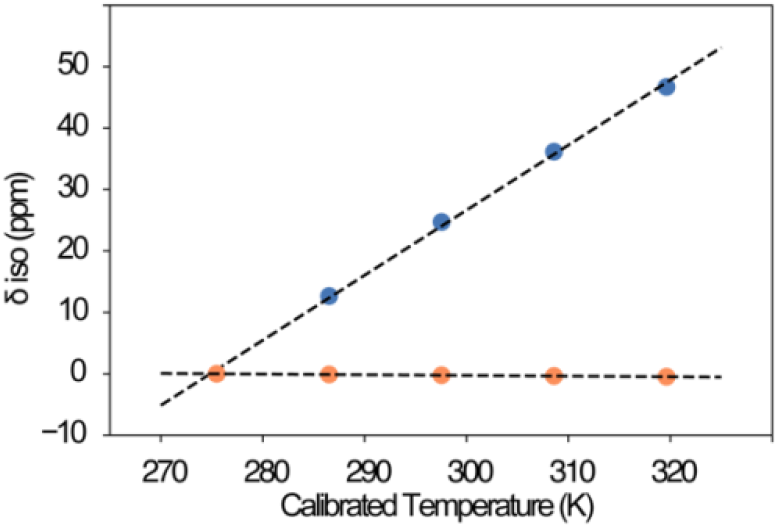
Comparison of the temperature dependency on the chemical shift of the H_6_ proton in TmDOTP (blue) and water proton (orange). Δδ_iso_ is the change in chemical shift relative to the shift at 275 K. All spectra were collected with spinning frequency at 5 kHz and gas flow rate at 1070 1/h. The dashed lines represent linear least squares fitting to the data: 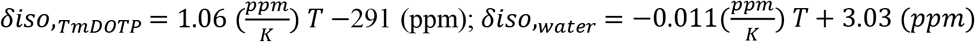. The error bars in both x and y dimensions for each data point are too small to be visualized. The error on slopes are dominated by the uncertainty from fitting. The ratio of slope and FWHM for TmDOTP and water are 0.7±0.3 and 0.09±0.01 respectively. The expansion of water chemical shift vs. calibrated temperature is shown in supplementary information (Figure S1).

### Spinning heating scales with the rotor frequency

MAS induced sample heating was measured and fit in a 3.2 mm E-free (Figure 4) and a 1.3 mm probe (Figure S2). Samples with just TmDOTP buffer and KcsA proteoliposome were used to demonstrate the negligible dependency of the MAS heating on sample. The spinning frequencies and corresponding sample temperatures of TmDOTP were fit to second-order polynomial function according to previous studies: 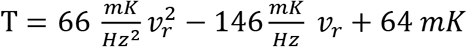, where = *ν_r_* = *ω_r_*/2π Notably, the H_6_ proton linewidth increased consistently with spinning frequency from 1636 Hz (at 2 kHz MAS) to 3663 Hz (at 16 kHz MAS). This increase inline width with MAS suggests that a heating gradient was caused by MAS. The line shape also grew more asymmetric with increasing MAS.

**Figure 2.**
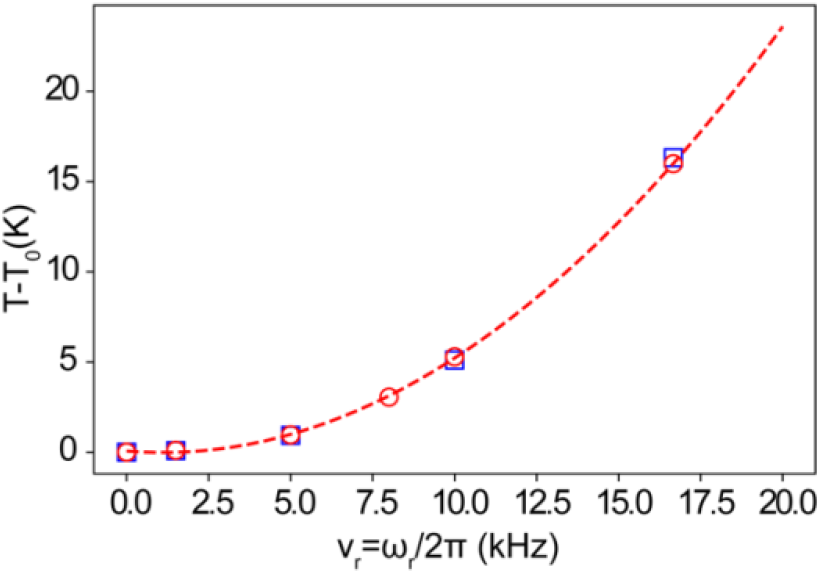
Sample temperature calculated from the H_6_ proton of TmDOTP as a function of spinning frequency. T_0_ is the temperature at zero spinning asymptote. TmDOTP buffer (red open circle) and KcsA proteoliposome samples (blue open square) were used here to demonstrate that MAS heating has little sample dependency. Data from the TmDOTP buffer were fit to second order polynomial function (red dash line): 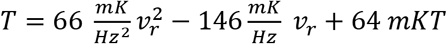 Data for 1.3 mm probe is shown in supplementary information (Figure S2).

### The influence of TmDOTP on hydrated proteoliposome sample

We compared the KcsA proeoliposome spectrum with and without 25 mM TmDOTP and observed no significant changes in chemical shifts or overall spectral quality (Figure 5A). KcsA, a pH activated potassium channel from *Streoptomyces lividans*, is used here since the marker peaks of the protein are sensitive to pH, temperature and potassium ion concentration changes^3,21^. This indicates that the paramagnetic nature of the TmDOTP complex has little to no effect on the properties of biological samples at the concentrations used. Marker peaks that belongs to the selectivity filter residues T74, T75 and V76 are specifically examined here (Figure 5B-D).

**Figure 3.**
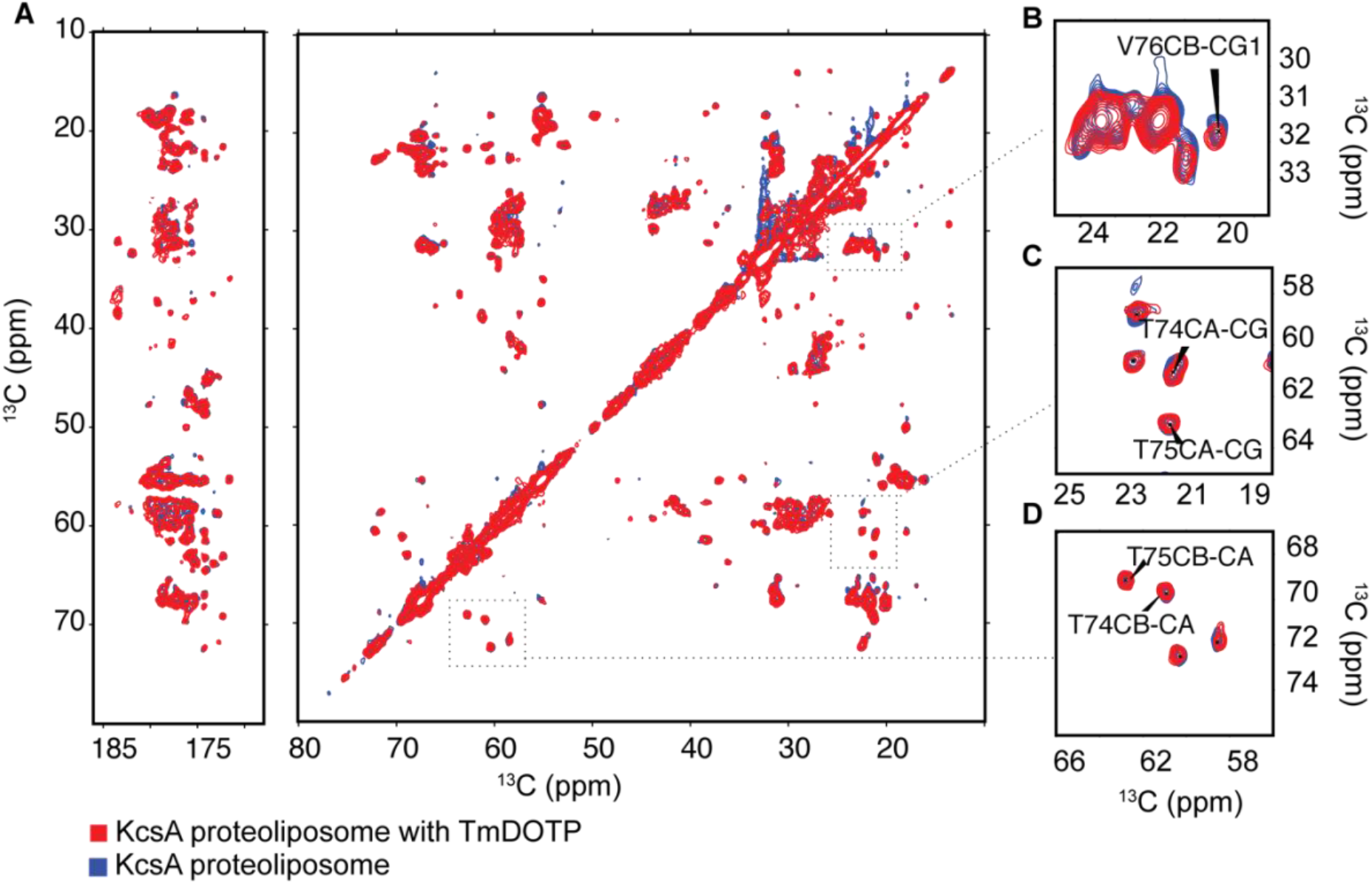
The effect of 25mM TmDOTP on KcsA proteoliposome. (A) Overlay of 2D ^13^C-^13^C correlation spectra of KcsA (blue) and KcsA with 25Mm TmDOTP (red) acquired in DOPE/DOPG (3:1) liposome at pH7.5. The regions of KcsA selectivity filter marker peaks are highlighted and shown in (B) V76 C β-Cγ (C) T74 Cα-Cγ and T75 Cα-Cγ (D) T74 Cβ-Cα and T75 Cβ-Cα. The data suggest no significant changes in protein structure and conformation state with the addition of 25mM TmDOTP.

### RF heating is linear with pulse power, pulse length and duty cycle

To examine the heating of a biological sample during RF irradiation, the pulse sequence shown in Figure 2 was applied to a KcsA proteoliposome sample and the temperature was monitored using the chemical shift of the H_6_ proton in TmDOTP. The target temperature was set at 275K on the VT control and the gas flow rate was 1070 L/hr. Continuous wave (CW) irradiation that resembles proton heteronuclear decoupling was applied here and the RF power, pulse duration (*τ*_1_), and duty cycle were varied respectively. *τ*_2_ was kept small (5 ms) here to limit sample cooling before acquisition. Figure 6 shows that heating is proportional to the RF power, duration of the pulse and duty cycle as discussed in the literatures^12,14^.

**Figure 4.**
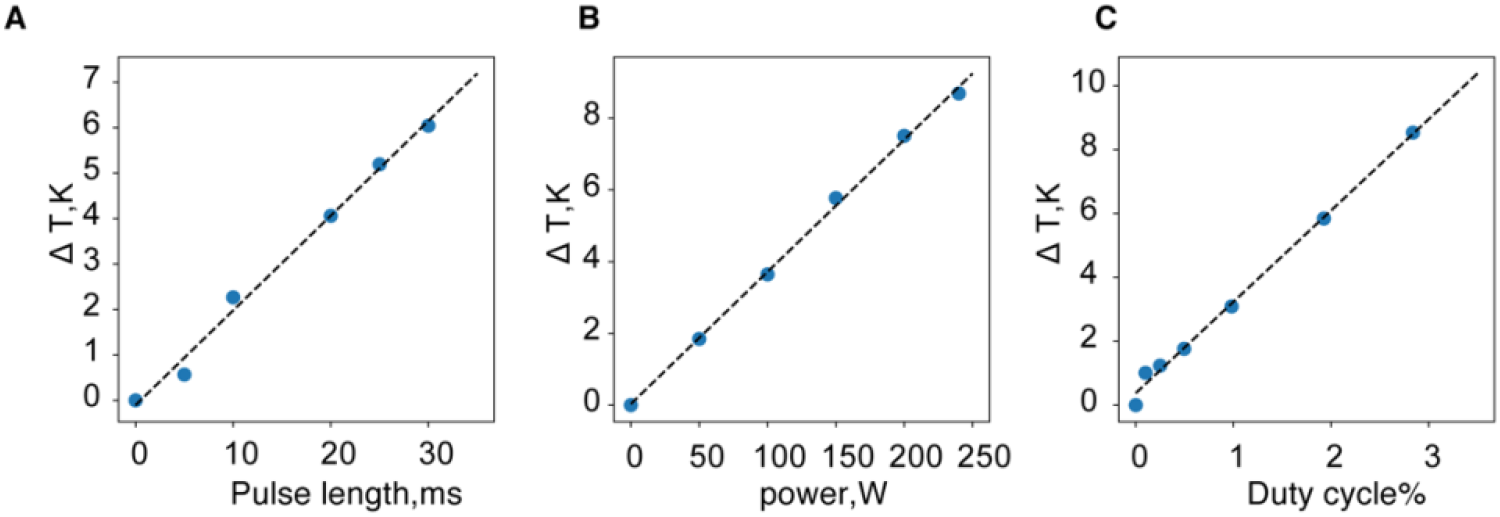
Plot of temperature increase calculated from TmDOTP chemical shift vs. (A) RF pulse length (field strength=91 kHz, duty cycle=2.8%). Data were fit using linear least square analysis (ΔT=0.208τ_1_-0.105). (B) CW power amplitude (*τ*_1_ = 30 *ms and duty cycle* = 2.8%). Data were fit using linear least square function (ΔT=0.037P+0.032), P stand for RF power (C) Duty cycle% (field strength=91 kHz and *τ*_1_ 30 *ms*). Data were fit using linear least square analysis (ΔT=2.860D+0.375). D stands for duty cycle%. All data were collected on a KcsA proteoliposome sample with 20 mM TmDOTP on 3.2 mm E-free prob at 900 MHz. The spinning frequency was 5 kHz and the target temperature was set at 275 K.

### Inequivalent RF heating on pellet vs. supernatant

One surprising finding from our RF irradiation study on the proteoliposome KcsA sample is that the H_6_ proton in TmDOTP peak splits into two components (denoted by peak 1 and peak 2) under RF irradiation on 3.2 mm E-free probe (Figure 7). The appearance of peak 2, which has a larger heating slope, only appeared under RF heating conditions, but not MAS (Figure S3). Moreover, the temperature reported by peak 2 matches with the one calculated from water proton chemical shift in the sample (Figure S4). However, the two distinct temperature populations are not resolved in water proton peaks possibly due to the broad linewidth (130 Hz). Therefore, we assigned the two peaks to TmDOTP in the pellet (peak 1) that sediments to the inner rotor wall due to the centrifugal forces generated by MAS and the peak 2 to the TmDOTP remaining in the center supernatant based on the agreement with the bulk water temperature measurement. The homogeneous linewidth calculated from T2 for the peak 1 and peak 2 are 815±39 Hz and 598±22 Hz respectively. The divergent temperatures indicated by TmDOTP peaks might arise from different cooling speed along the radial axis of the rotor and the distinct heat capacity of water and proteoliposome. The data collected on a 1.3 mm solenoid probe is shown in Figure S5. At the same field strength, the uppermost temperature of the sample is consistently higher on 1.3 mm probe than 3.2 mm E-free probe and the heating gradient is continuous rather than peak splitting. The determined heating gradient can be as large as 18 K at the field strength of 90 kHz using the 1.3 mm probe. The difference in line shape between 3.2mm E-Free and 1.3mm solenoid probes is due to the probe design and heating/cooling mechanism.

**Figure 7.**
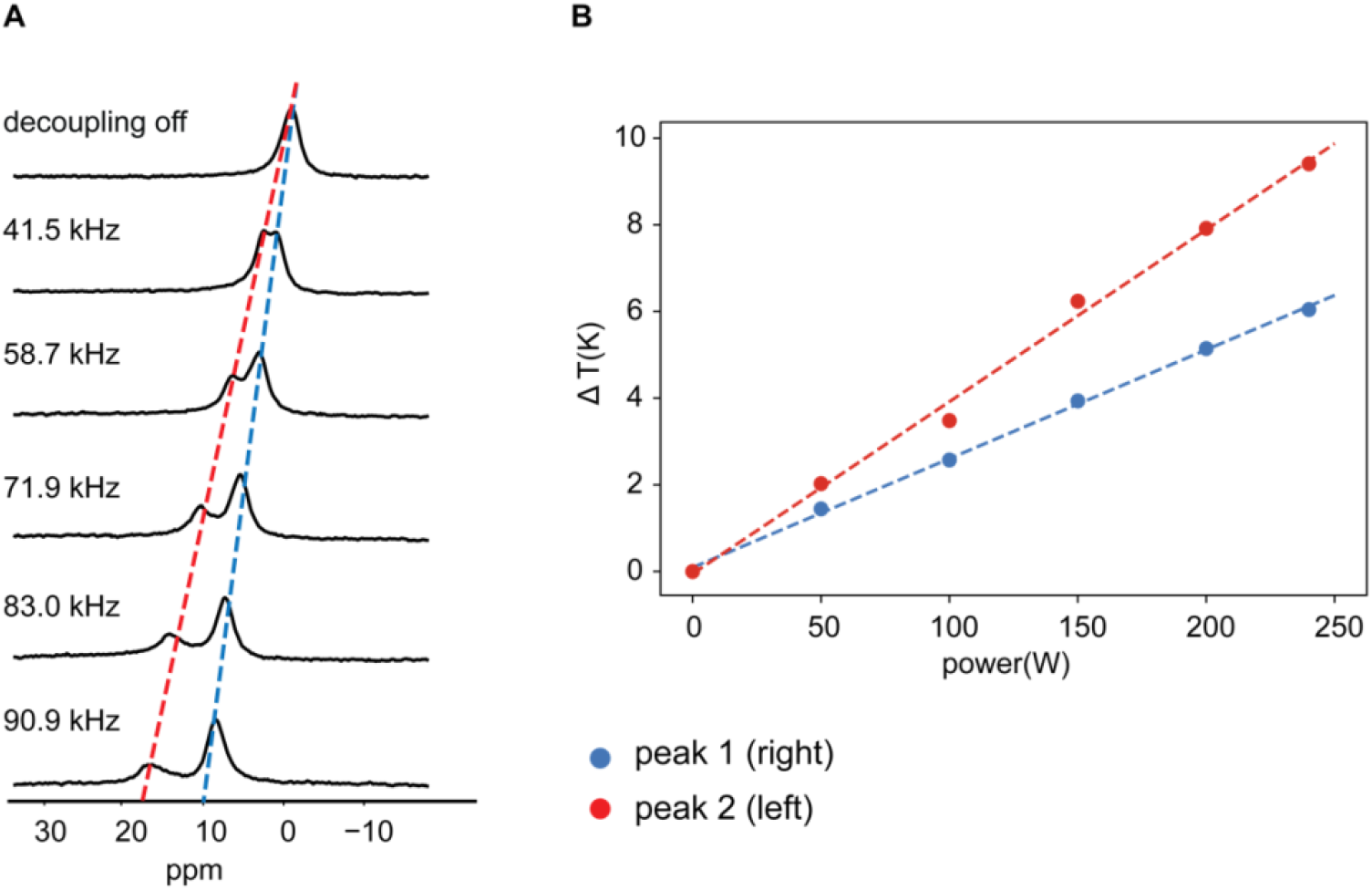
(A) The H_6_ NMR spectra of the proteoliposome sample with 20 mM TmDOTP at neutral pH during different RF frequencies. Target temperature was set to 275 K. The chemical shift of H_6_ was set to 0 ppm when the decoupling pulse was off. This adjustment is for the convenience in temperature reading, since TmDOTP has a slope of nearly 1 ppm/K. (B) Plot of RF power vs. sample temperature changes reported by the peak 1 and peak 2 from TmDOTP proton chemical shifts. All the data were collected on the 3.2 mm E-free probe at 900 MHz.

### Application and significance

Owing to the fast relaxation rate and minimal perturbation on biological sample properties, TmDOTP can be incorporated into SSNMR samples to monitor real time temperature throughout an experiment. This may be crucial for samples and measurements that are sensitive to temperature changes, such as R_1ρ_^26^. Here, we demonstrate the temperature mapping of a ^13^C-^13^C dipolar assisted rotational resonance (DARR) experiment using 20 mM TmDOTP in KcsA proteoliposome sample. The experiment was carried out at Bruker 700 MHz equipped with a 3.2 mm E-Free probe under 12.5 kHz MAS. The multi-receiver feature on AVANCE NEO enabled an immediate H_6_ chemical shift measurement following every carbon acquisition. Figure S6 shows the temperature of KcsA sample increased about 0.5 K through the experiment caused by proton high power (90 kHz) proton decoupling during the increasing evolution time t1. In addition, our data demonstrate that the heating from MAS and RF radiation are not additive. In order to obtain the precise temperature during an experiment, it is necessary to include a real time thermometer, such as TmDOTP, rather than simple extrapolation (Figure S7).

## Conclusions

With a linear temperature dependency and large thermal resolution, we demonstrate that TmDOTP is an excellent internal thermometer for solid state NMR experiments on biological samples. The distinct proton chemical shift and short T1 enable an instant reading of the precise temperature in a biological sample. Comparing with common thermometer molecules that employed in SSNMR, such as ^207^Pb in Pb(NO_3_)_2_, ^119^Sn in Sm_2_Sn_2_O_7_, and KBr, TmDOTP stands out in its low toxicity and nearly negligible perturbations on sample properties. Moreover, the proton detection eliminates the change in probe configuration and enables real time temperature reading and heat distribution measurement throughout an experiment. In addition, we observed two discontinuous temperature population in KcsA proteoliposome sample induced by RF irradiation. The two peaks were assigned to the pellet at the inner rotor wall and the center supernatant that result from MAS centrifugation. Finally, the discrepancy between the real temperature of decoupling while spinning and the extrapolated value from MAS and RF heating fitting curve shows the necessity and importance of obtaining real time temperature.

## Acknowledgments

The NMR data was collected at the New York Structural Biology Center (NYSBC) with support from the Center on Macromolecular Dynamics by NMR Spectroscopy (CoMD/NMR) a Biomedical Technology Research Resource (BTRR) supported by U.S. National Institutes of Health (NIH) through grant number: P41 GM118302. The NYSBC is also enabled by a grant from the Empire State Division of Science Technology and Innovation and Office of Research Infrastructure Programs/NIH Facility Improvement Grant CO6RR015495. A.E.M. is a member of the NYSBC. This work was supported by NIH Grant R01 GM088724 (to A.E.M).

