## Supplementary figures for "TmDOTP : An NMR- based Thermometer for Magic Angle Spinning NMR Experiments"

**Supporting Information**

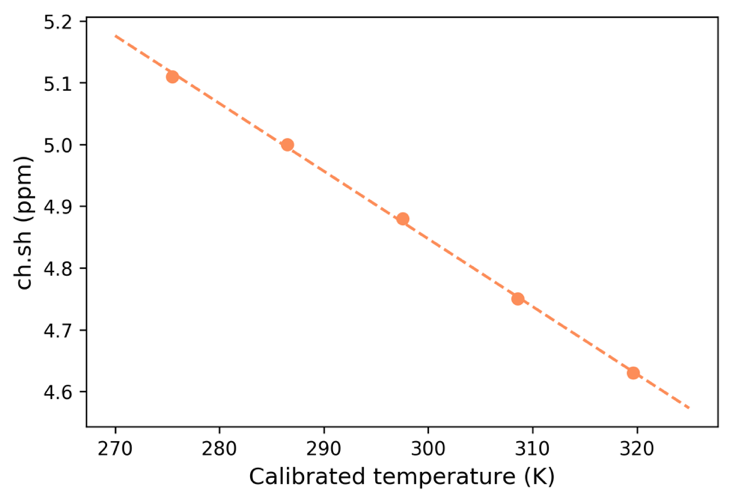

**Figure S1**. Plot of water proton chemical shift vs. temperature. All data were collected on 900MHz and 3.2mm E-free probe.

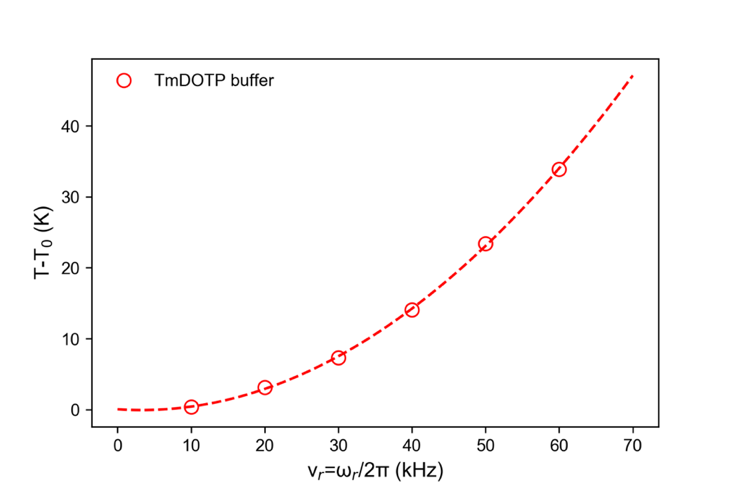

**Figure S2.** Sample temperature calculated from ^1^H TmDOTP as a function of spinning frequency on Bruker 900 MHz and 1.3 mm probe. T_0_ is the temperature at zero spinning asymptote. Data from the TmDOTP buffer were fit to second order polynomial function:$T=0.011\omega_{r}^{2}-0.068\omega_{r}+0.038$.

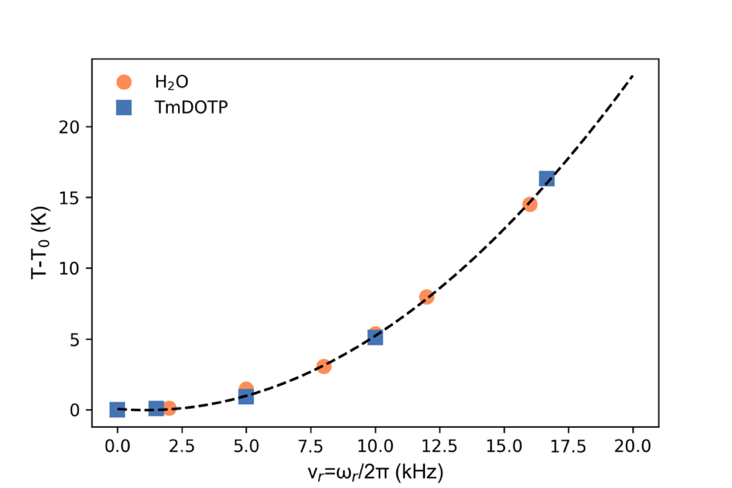

**Figure S3** Plot of sample temperature calculated from ^1^H TmDOTP and water proton in the KcsA proteoliposome sample as a function of spinning frequency on Bruker 900 MHz and 3.2 mm probe. T_0_ is the temperature at zero spinning asymptote.

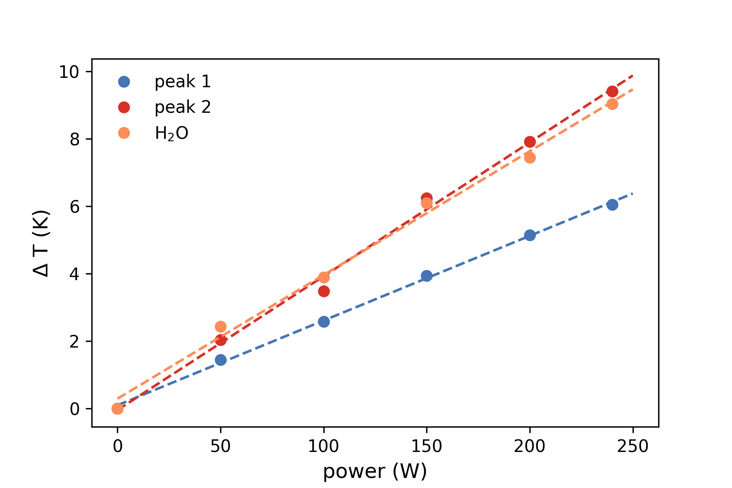

**Figure S4.** The plot of RF power vs. sample temperature change calculated using the H_6_ proton in TmDOTP and water proton chemical shift in KcsA proteoliposome sample at pH 7.5. The MAS was 5 kHz and temperature was 275 K ($\tau_{1}=30 ms, \tau_{2}=500 ms, recycle delay=1 s$).

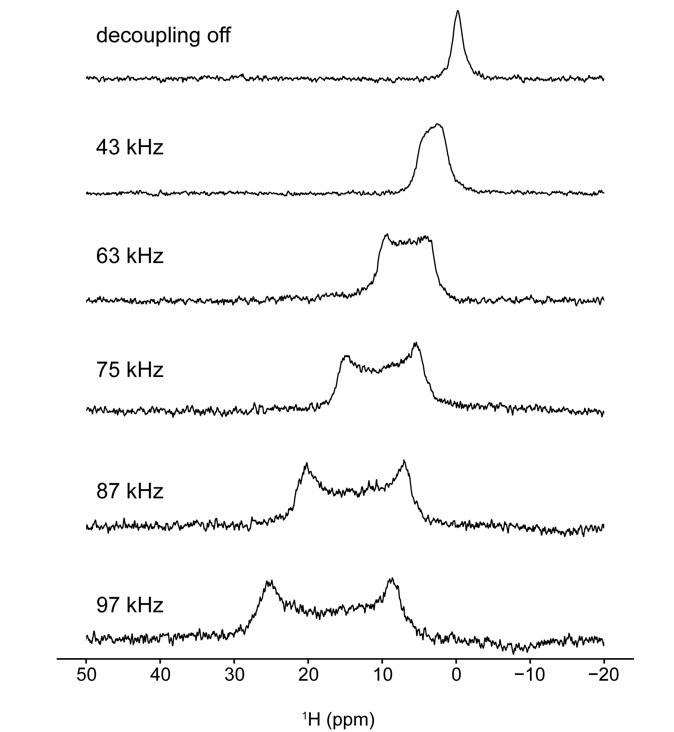

**Figure S5.** TmDOTP spectra obtained at different RF powers on 1.3 mm HCN Probe with MAS at 5 kHz. The TmDTOP H_6_ proton spectra of the proteoliposome sample were collected at 275 K. The chemical shift of H_6_ was set at 0 ppm when the decoupling pulse was off. This adjustment is for the convenience in reading the temperature changes, since the H_6_ proton in TmDOTP has a slope of near 1 ppm/K. RF heating on solenoid probe shows significant large heating gradients at high RF power.

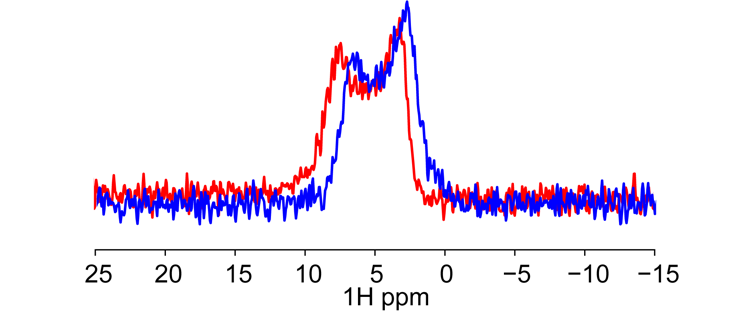

**Figure S6**. Overlay of the ^1^H spectrum of the H_6_ proton in TmDOTP at the 1^st^ (blue) and the 512 (red) slice in indirect dimension of the ^13^C-^13^C DARR experiment (number of scans=16). High power decoupling (90 kHz) was applied during evolution time t1 and the increment delay was 28.39 μs. The experiment was carried out on a Bruker 700 MHz at 12.5 kHz MAS. The temperature was set at 275 K on VT controller and the VT gas flow was 2000 l/h. The KcsA proteoliposome sample was prepared as described in experiment and method section.

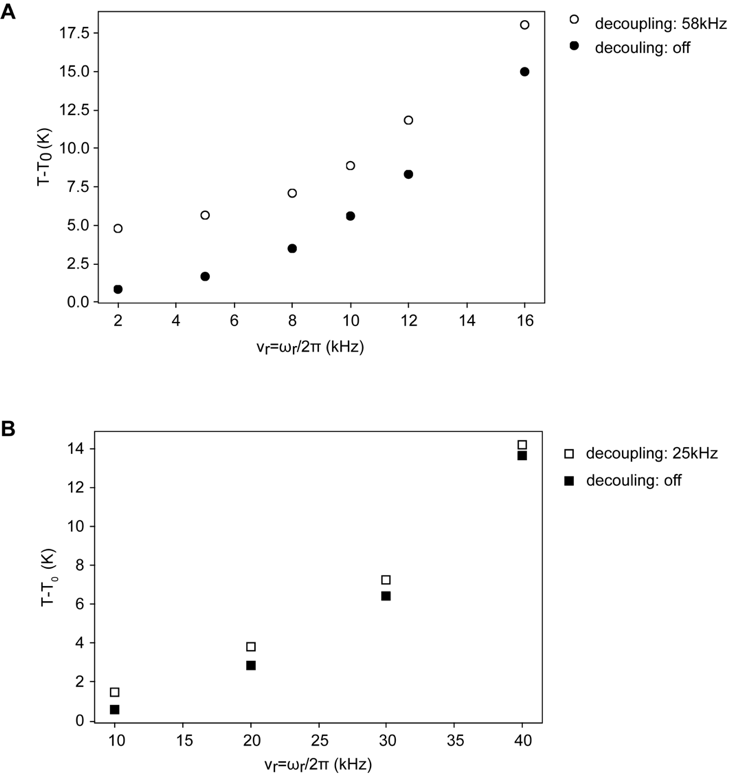

**Figure S7**. Combined heating effects from spinning and RF irradiation (A) Sample temperature calculated from the H_6_ proton of TmDOTP as a function of spinning frequency with decoupling pulse on(open circle) and off (solid circle) on 3.2 mm E-free probe. (B) Sample temperature calculated from the H_6_ proton of TmDOTP as a function of spinning frequency with decoupling pulse on(open square) and off (solid square) on 1.3 mm solenoid probe. T_0_ is the temperature at zero spinning asymptote. Data were collected on Bruker 900 MHz at 275 K.
